# CaBLAM for chiropraxis in cryoEM, UnDowser to rethink “waters”, and NGL Viewer to recapture online 3D graphics in MolProbity validation

**DOI:** 10.1101/795161

**Authors:** Michael G. Prisant, Christopher J. Williams, Vincent B. Chen, Jane S. Richardson, David C. Richardson

**Affiliations:** Department of Biochemistry, Duke University Medical Center, Durham NC 27710

**Keywords:** Structure validation, backbone conformation, overfitting, water analysis, server security, NGL Viewer, Ramachandran restraints, hydrogen bonding

## Abstract

The MolProbity web service provides macromolecular model validation to help correct local errors, for the structural biology community worldwide. Here we highlight new validation features, and also describe how we are fighting back against recent outside changes that degrade or endanger that mission. Sophisticated hacking of the MolProbity server has required continual monitoring and various security measures short of restricting user access. Refinement software now increasingly restrains validation criteria in order to supplement the sparser experimental data at 3-4Å resolutions typical of modern cryoEM. But unfortunately the broad density allows optimization without fixing underlying problems, which means these structures often score much better than they really are. CaBLAM, our first new tool designed for this regime, was described in the previous Tools issue, and here we demonstrate its effectiveness in diagnosing local errors even when other validation outliers have been artificially removed. The deprecation of Java applets now prevents KiNG interactive online display of outliers on the 3D model during a MolProbity run, but that important functionality is now recaptured with a modified version of the Javascript NGL Viewer. Other new changes are more straightforwardly good. We are moving to the Neo4j database (graphical rather than relational), and will soon have cleaner as well as much larger reference datasets. In addition to several minor new features, we have developed a tool called UnDowser that analyzes the properties and context of modeled but clashing HOH “waters” to diagnose what they might actually represent. A dozen distinct scenarios are illustrated and described.

## I. Introduction

MolProbity does macromolecular model validation across a suite of criteria, for X-ray, neutron, NMR, computational, and now cryoEM models (Davis et al. 2004; Davis et al. 2007; Chen et al. 2010; Williams et al. 2018a). It is a current widely-used system (the papers have over 12,000 total citations), within a 30-year history, where advances in validation methods have been prompted by advances in structural biology capabilities or by crises of trust.

The field of structure validation began around 1990, when crystal freezing, synchrotron data, and more powerful refinement methods such as simulated annealing in XPLOR (Brunger 1988) lowered R-factors, invalidating prior rules of thumb for when a structure had reached acceptable accuracy. That led to several serious mis-tracings and high-profile retractions such as for RuBisCo (Knight et al. 1989) H-ras p21 (Pai et al. 1990) and HIV protease (Wlodawer et al. 1989), which in turn prompted the development of Rfree (Brunger 1992), Oops (Jones et al. 1991), ProCheck (Laskowski et al. 1993), WhatCheck (Hooft et al. 1996), and other validation systems.

By about 2000, more full-system automation had opened crystallography to non-experts, and was both exploited and advanced by Structural Genomics centers. Distrust of black-box crystallography prompted community demand for required validation in the customary “Table 1” of structure papers. For our research group, the driving factor was the failure of early protein *de novo* design to produce well-packed interiors and avoid molten globules (Richardson et al. 1992; Richardson & Richardson 2013). That problem led us to develop optimized H addition & all-atom contact analysis for quantification of packing quality (Word et al. 1999a; 1999b). The contact analysis turned out also to be a powerful new tool for model validation, and its local and directional nature guides correction of the flagged errors. That process was extensively tested in production use (Arendall et al. 2005) and has helped structural biologists improve their models ever since (see 1b), up to about 2.5Å resolution.

One investigator’s set of 11 fraudulent structures (Janssen et al. 2007; Borell 2009) led to the worldwide PDB’s Validation Task Forces for community organized standards (Read et al. 2011; Montelione et al. 2013), validation on deposition including reports for referees (Gore et al. 2012), and many further developments.

Just recently, the cryoEM revolution has produced an urgent need for better validation at 2.5 to 4Å resolution, prompting new methods and tools including EMRinger (Barad et al. 2015), Qscore (Pintile & Chiu 2018), and the current & planned changes in MolProbity described here. These respond both to excellent things, such as high-resolution cryoEM and the expansion in use of ensemble structures, and to bad things, such as targeted website attacks and the fact that at 2.5 to 4Å the broad density necessitates modeling and refinement which directly or indirectly restrains most current validation criteria and makes the structures score better than they really are.

## II. Results

### II.1 Indicators of progress

In the previous Tools issue (Williams et al. 2018a) we described the epidemic overuse of unfavorable and very rare *cis*-nonPro peptides; only 1 in 3000 residues are genuine, usually functional occurrences (Williams et al. 2018b). We showed that since flagging of *cis* and twisted peptides has been added to MolProbity, Phenix, and Coot, there seemed to be improvement in that problem It is now clear that overuse of *cis*-nonPro has indeed gone back down nearly to pre-epidemic levels (Figure 1a).

**Figure 1.**
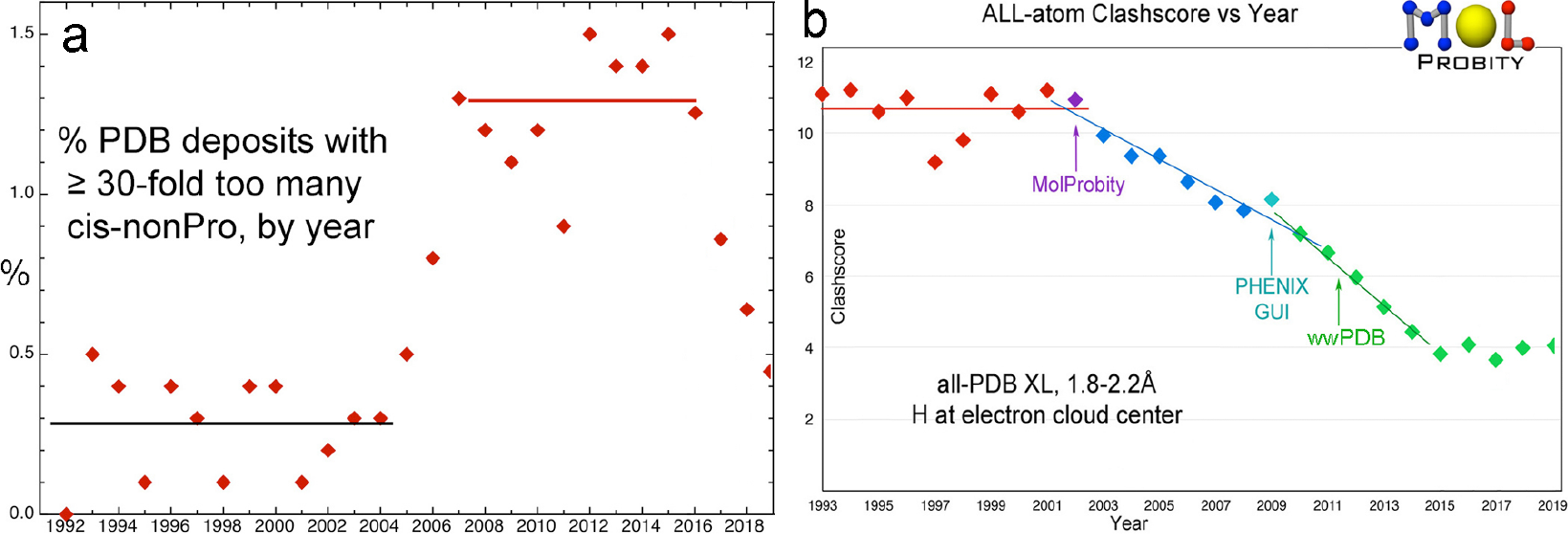
Timeline of MolProbity validation metrics. a) Overuse of very rare *cis*-nonPro peptides in deposits to the wwPDB (Berman et al. 2003), showing the abrupt rise around 2006 (Croll 2015; Williams & Richardson 2015) and the now successful return to pre-epidemic levels. b) Mid-resolution (1.8-2.2Å} clashscores (number of non-H-bond steric overlaps ≥0.4Å per 1000 atoms), steadily decreasing since 2002 and now level at about 4.

Mid-resolution clashscores (number of non-H-bond steric overlaps ≥0.4A per thousand atoms) for worldwide PDB depositions have been steadily decreasing since the 2002 introduction of all-atom contact analysis by MolProbity. That metric has now leveled off at a reasonably low level of about 4 on average (Figure 1b). By definition clashscores cannot go below zero, and a clashscore less than about 2 is often a result of overfitting, since 2.7 is the average score in the best parts of the quality-filtered, high-resolution reference data.

Another very positive development is that in addition to increasingly thorough integration of model validation between the MolProbity web service and the validation GUIs in Phenix (Liebschner et al. 2019), MolProbity validation is now also available in the CCP4 system (Winn et al. 2011).

### II.2 New features: UnDowser to diagnose non-water HOHs

At high to mid resolutions better than about 2.5Å, density peaks can be seen for individual water molecules bound dynamically but reproducibly to favorable sites where they H-bond to protein, nucleic acid, or ligands. On an initial model, these will be positive difference peaks, into which waters are then fit either manually or automatically. However, it is well known that not all such peaks actually represent waters. The clashes of our all-atom contact analysis provide powerful criteria especially useful for distinguishing the wide variety of cases that are something other than water (Headd et al. 2013), and applicable even to low-coordination sites at the molecular surface. A new feature in MolProbity called UnDowser now produces a table to show such diagnosis, to guide the user in making these decisions. The table includes all clashing HOHs, sorted in approximate order of severity (by Sum(overlap −0.2Å). Type and charge of the atom(s) with which they clash are noted, and probable diagnoses of the underlying problem are given. Examples are shown and discussed here below for about a dozen distinct scenarios, to aid user comparison with their own cases.

An HOH that clashes with two or more atoms of the same polarity, and with no nonpolars or opposite polars, is almost certainly an ion. If all clashes or H-bonds are with negative atoms, then the HOH is a **positive ion**; if all interacting atoms are positive, then the HOH is a **negative ion**. Such interactions show up graphically as hotpink clash spikes inside the green dots of an H-bond. Occasionally, full coordination is seen, as shown in Figure 2a for the positive ion modeled as HOH 606 in the 6hhm sulfatase (Reisky et al. 2019), with two clashes to oxygen atoms. That HOH has 6 ligands -- 3 backbone CO, 2 Thr Oγ, and a water -- in closely octahedral geometry. Once an ion has been explicitly modeled, coordination-geometry tools (Echols et al. 2014; Zheng et al. 2017) can identify the most probable ion species, in this case Na+.

**Figure 2.**
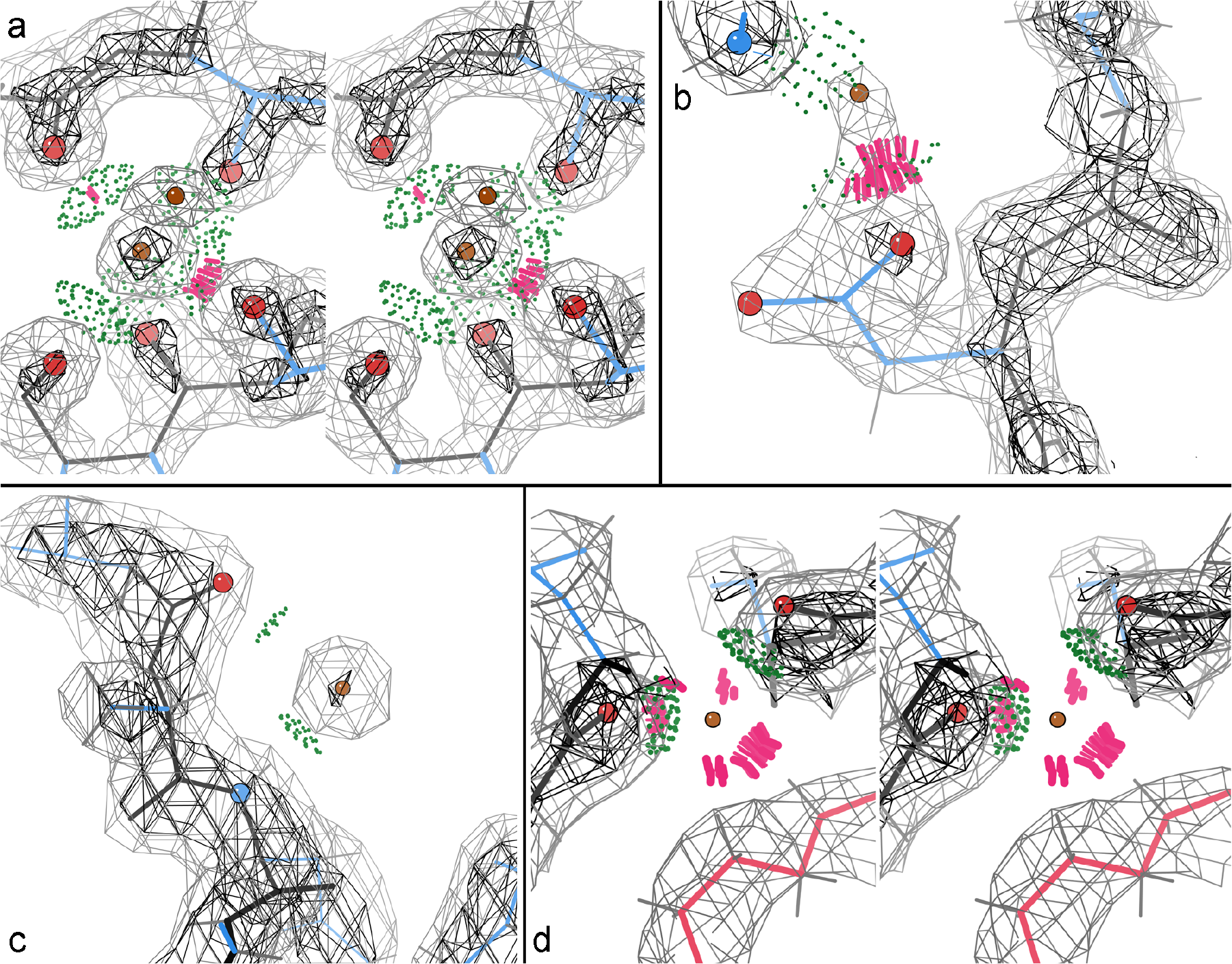
UnDowser diagnosis of HOH “waters”. a) Stereo of a positive ion modeled as HOH 606 in 6hhm (Reisky 2019), with 6 near-octahedral oxygen ligands. Lenses of green all-atom contact dots show donor-acceptor overlaps, but 2 distances are so short they also have serious clashes (clusters of hotpink spikes). b) A confusing HOH in 6hhm with one clash to an Asp carboxyl O, but also a good H-bond to an Arg guanidinium. c) A genuine water, with strong, round density and 2 H-bonds with atoms of opposite partial charge. HOH 414 in 6a4v (Hwang unpublished). d) Stereo of HOH 504 in 5onu (Li 2018)with one polar and 4 nonpolar clashes. There is no electron density even at zero, suggesting deletion.

If there is only one HOH-to-polar clash the diagnosis can be complex and depends on many factors of the context, such as clashes or close interactions with nonpolar or wrong-charge atoms, and relative density strength, B-factors, and shape of the groups involved. Clearly diagnosable cases that include a single HOH-polar clash show up in three later figures, and an ambiguous case is discussed here. A **partially-occupied +ion at a carboxyl O** is relatively common, where the interacting O usually shifts somewhat when unbound. At high resolution a clashing water might be modeled, but at mid resolution this may just produce a strangely shaped sidechain density with an extra lobe. Figure 2b shows a strange triangular density shape with an extra lobe for Asp 388 of 6hhm at 1.23Å resolution. The HOH modeled in that lobe cannot be either a water or an alternate conformation of the Asp (the lobe is too far from the Cα and too close to the Oδ). An ion is however rendered unlikely by the good H-bond to Arg 341 Hh1. In this confusing case the HOH should perhaps be deleted.

In contrast, **genuine water**s often H-bond to atoms of opposite charge, since their tetrahedral coordination includes two donors and two acceptors. Figure 2c shows a real water in the all-helix viral ORF of 6a4v, with good 2.2Å density and weak but reasonable H-bonds to backbone NH and CO. A non-clashing HOH with a well-separated, round density peak at good H-bonding distance and angle from one or more polar atoms is nearly always a genuine water.

An extreme, unambiguous case is when after refinement there is extremely weak or absent density at the HOH position. Such a **“water” should be deleted**, as for the case shown in Figure 2d of an HOH at 2.22Å in the OmpU trimer of 5onu (Li et al. 2018). It superficially looks rather like a coordinated ion, but 4 of the clashes are to nonpolar atoms and the HOH peak has no density even at zero contour level. At present, the user must diagnose this, since inside the MolProbity website UnDowser does not have access to density maps (a future implementation inside Phenix will use information from electron density).

A similar-appearing case of many large clashes, including nonpolars with gorgeous density, is seen for HOH 51 and others in 3azd at 0.98Å. Apparently this rarely-seen problem is that the data were detwinned, lowering the information content enough that the 2mFo-DFc map is very nearly a calculated map which thus slavishly follows even incorrectly modeled features. Comparison of B-factors between the HOH and surrounding atoms is often inconclusive because of B-factor restraints and partial-occupancy waters, but when extreme mismatches occur they offer a reason to look. For the 5onu case above, the ghost water has a B of 137 while the surrounding protein atoms average 45. On the other hand, an HOH B-factor much lower than surrounding atoms or than well-ordered backbone O, that suggests a **heavier atom than O** even if it does not clash with anything.

If a water looks good but clashing atoms have higher B-factors, weaker density, or poorly fit density, then try fitting a **different conformation for the clashing group** that preferably preferably H-bonds to the water rather than clashes. This often happens at lysine Nz, as shown in Figure 3a for the strong HOH B283 at 2.0Å in 6aht (Hayashi et al. 2018), which clashes with the weak end of Lys B106. Figure 3b shows a rebuilt, H-bonding lysine rotamer. The clashing atom can be at fault even with good density, if it is in a **backward-fit sidechain amide or histidine**, as shown in Figure 3c for HOH 1033 clashing with Cδ2 of the backward-fit His 73 of 1bkr at 1.1Å (Banuelos et al. 1998).

**Figure 3.**
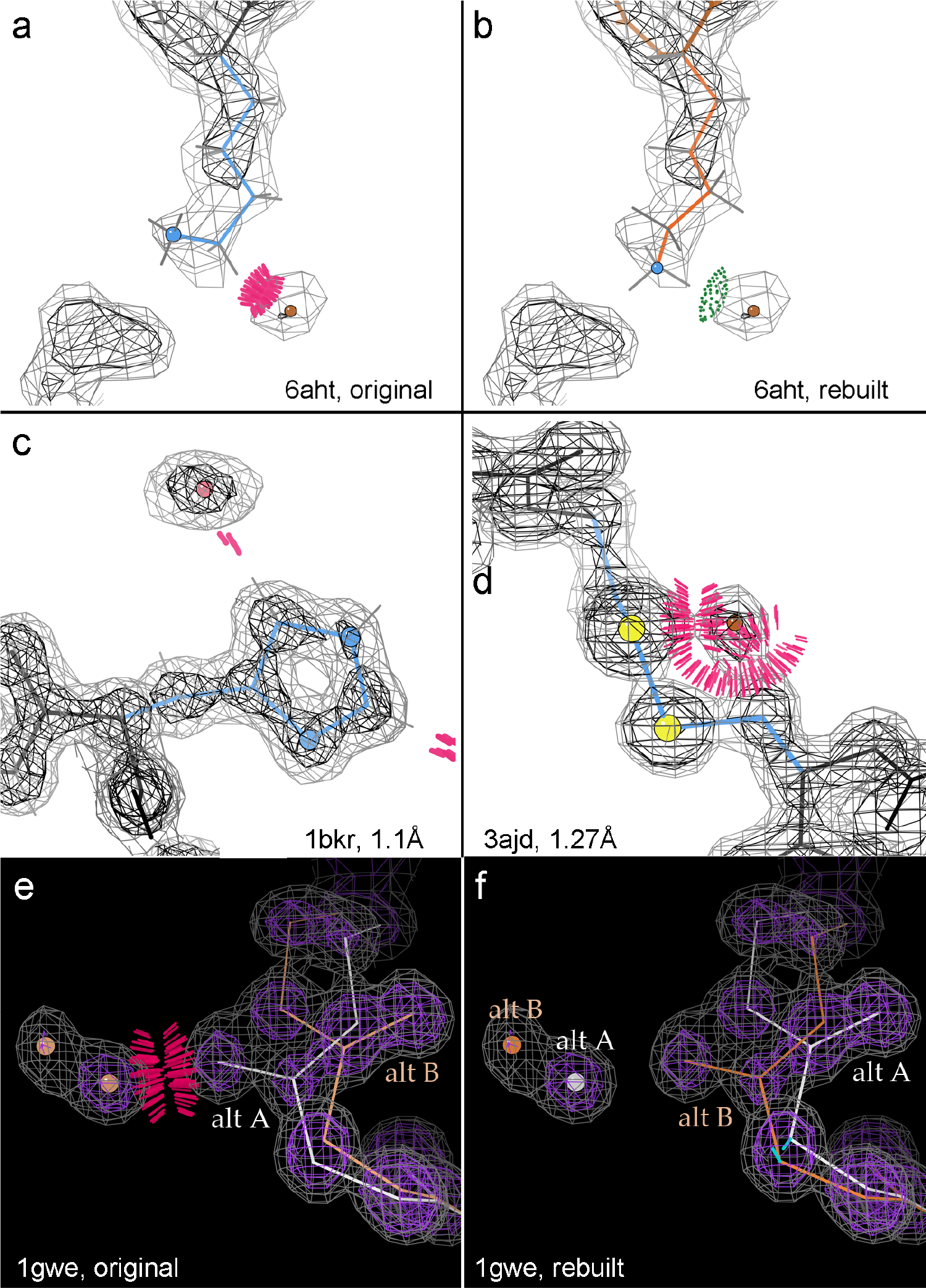
HOH “waters”, context interpreted: a) HOH B283 clashes with Lys methylene at 2.0Å in 6aht (Hayashi et al. 2018). b) Rebuilt Lys H-bonds instead. c) Clash of HOH 1033 with Cd2 of the backward-fit His 73 of 1bkr at 1.1Å (Banuelos et al. 1998). d) Shows a “water” that is really an alternate conformation of the 91-186 disulfide bond in 3ajd at 1.27Å (Kuratani et al. 2010). e) A full occupancy HOH pair clashing with mislabeled alternates of Leu 105 at 0.88Å in 1gwe (Murshudov et al. 2002). f) Reassignment of higher-density Leu and water alternates as altA at 60% occupancy.

At high resolution, an HOH that clashes with a non-polar atom (especially if along a sidechain) is often the **next atom in an unmodeled alternate conformation**. Figure 8 of Richardson et al. 2018a shows two HOHs clashing with the Cβ methylene of Asp 9 in the Zn protease of 1eb6 at 1.0Å (Macauley et al. 2002), cleanly corrected with a small “backrub” adjustment (Davis 2006) and the most common Asp rotamer. A similar situation is shown in Figure 3d for an HOH where the 186 S should be, for an unmodeled alternate conformation of 91-186 disulfide bond in 3ajd at 1.27Å (Kuratani et al. 2010). A partially reduced disulfide, common from radiation damage during data collection, might also have one or both SHs incorrectly modeled as HOH.

On the other hand, an HOH that clashes with an already-modeled alternate conformation protein atom most likely is a **real water that needs an occupancy <1.0, or** sometimes the **altA, altB naming is incorrect**. Both those situations occur in Figure 3e for the full-occupancy HOH pair and Leu 105 mislabeled alternates at 0.88Å in the 1gwe bacterial catalase (Murshudov et al. 2002). Figure 3f shows a rebuild with the higher-density Leu and water alternates as altA at 60% occupancy and the lower-density ones at 40%. Each alternate-conformation model is now clash-free and forms a water-backbone H-bond.

Two or more HOHs that clash with one another usually just **need compatible partial occupancies**<1.0. In fact, this is so common that the standard clashscore in MolProbity does not calculate or include clashes between waters; that is implemented only within UnDowser. However, two or more clashing HOHs may well represent an **unmodeled ligand** such as sulfate, GOL, or PEG. Figure 4a shows a set of 3 very badly clashing HOHs at 1.77Å in the 1LpL CAP-Gly domain (Li et al. 2002), which H-bond with Lys 175 and Arg 212. Figure 4b shows this density rebuilt as a very convincing SO4 in 1tov (Arendall et el. 2005).

**Figure 4.**
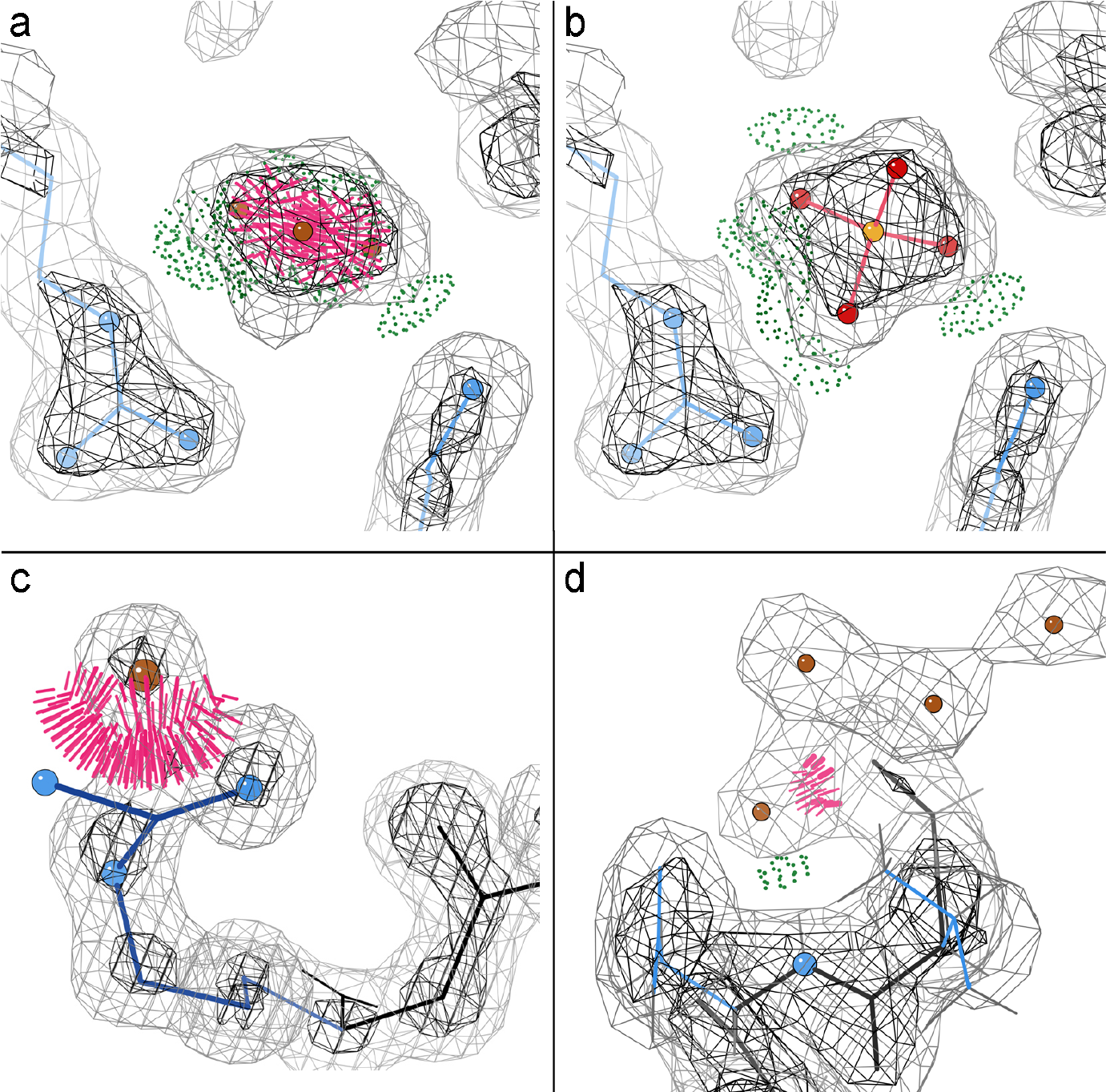
Sets of clashing HOH “waters” suggest reinterpretations. a) Three very badly clashing HOHs at 1.77Å in 1LpL (Li et al. 2002), which H-bond with Lys 175 and Arg 212, in b) reinterpreted as SO4 in 1tov (Arendall et el. 2005). c) A prematurely fit HOH A2090 forces the Arg A59 guanidinium out of density in 1qLw at 1.09Å (Bourne et al. 2000). d) Four HOH loosely fit into connected density, one clashing with N of the first-modeled residue in the 2.22Å OmpU of 5onu, suggest an additional, earlier helical turn of backbone.

A quite disruptive local problem is an HOH fit into **density that actually belongs to some other atom** which is thus kept out of its correct position, most often in a sidechain. This happens when the initial model (either molecular-replacement or *ab initio*) has the wrong rotamer. That produces a positive difference peak where the displaced atom should have been, and then automated water picking places an HOH there. Refinement cannot recover from this type of mistake, except for a procedure that deletes all waters before a rotamer-rebuilding step. However, an informed look at severely clashing, apparently strong “waters” can diagnose corrections on the basis of short distance and connecting density to sidechain atom(s) and poor fit for the sidechain. A stunningly obvious high-resolution example is shown in Figure 4c where HOH A2090 displaces an Arg A59 guanidinium N atom in the novel esterase of 1qLw at 1.09Å (Bourne et al. 2000), to serve as a guide for recognizing such patterns even at lower resolutions.

For backbone atoms, incorrect HOH mimicry is apt to be at a **non-terminal chain end that should be extended**. As an example, Figure 4d shows four HOH loosely fit into connected density, one clashing with the backbone N of Gly 32, the first-modeled residue in the 2.22Å OmpU of 5onu. That density almost certainly represents an additional, earlier turn of backbone, but the clashing HOH has displaced the carbonyl O atom of peptide 31-32.

For clashing waters, as well as for other validation outliers, the aim is to fix cases that are wrong in ways apt to be both significant and stable to further refinement: conformations in the wrong local minimum or groups that change existence or identity. The ability for clashes to flag meaningful errors of HOH identity depends on a good balance in the refinement target between steric overlap and density-fit terms. If sterics are over-weighted, there will be clashes only for the very worst cases. If density fit is over-weighted, then sometimes, at mid resolution and especially for cryoEM, there are many small water clashes spread throughout the structure, as seen for instance in the excellent 2.2Å 5a1a β-galactosidase structure (Bartesaghi et al. 2015). Most of these are not worth trying to fix individually. It might, however, be worth looking only at clashes >0.5Å (dark pink in the MolProbity multi-chart), to decrease the noise and find the more important problems.

This new water diagnosis, especially when it finds ions or ligands, can be of biological importance and is always an issue for computations based on the structure. We hope the UnDowser tool will make such choices much easier, even in this initial form, and its diagnostic criteria should continue to improve with experience in large-scale use.

### II.3 Security issues

For the last several years, our main MolProbity web service has come under specifically tailored, sophisticated web hacking attacks, serious enough to threaten continuing existence of the site and forcing intensive defense efforts. Here we give a brief history of the nature of these security threats, describe our defense strategies to keep this widely used community resource running, and note specific changes that may be visible to our users.

The foundational design of MolProbity occurred in a very different early internet era when one did not expect a strictly research service to be a hacking target. Since 2017, attacks on Molprobity have escalated to include thoroughly researched, specifically targeted hacks, presumably because it is located inside the network of a major medical center whose walls the attackers hope to breach.

Since its initial deployment, MolProbity has grown enormously in an organic fashion, with contributions from many authors. It is not a monolithic program but rather an ecosystem of diverse tools in many programming languages: C, C++, Java, Bash script, Perl, and more recently Python and Javascript. These executable tools are presented to the user through a dynamic, hand-coded web framework written in PHP. Users upload files for active analysis on the site and download the results. All of the code is open source, on GitHub. The PHP Query String Parameters that drive specific program execution are visible in the web URL request. Session management is file-directory based, with session state dynamically determined by session directory content. Those directories are also visible to an outside user, given modest reverse-engineering. Finally, our email bug report system is susceptible to constant spamming. In short, MolProbity is too large and complex for us to rewrite completely, but is intrinsically vulnerable in our current internet environment.

Given these realities, our approach has been a patchwork of partial changes to the system, combined with constant monitoring.

Changes to the system include: 1) The actual MolProbity server machine is now housed outside the lab on a private, secured subnet. So we do not have direct physical access to the machine, and thus restoration of service in case of outage is no longer fully under our control. If the main site is down, try the mirror site at molprobity.manchester.ac.uk, but note that it may serve an older version. 2) We have limited the variety of uploads or fetches, and filter all uploads for Trojan scripts. Users must now begin with a coordinate file and cannot upload maps or kinemages for online display. We also check for well-formed PDB files, both at input and during processing, with the useful by-product of more helpful error messages for the commonest problems. 3) We have made modest changes to obstruct wide-open access to MolProbity’s data directories, placing our administrative tools behind Apache basic password protection. 4) We are in the process of migrating to a different bug-report mechanism less susceptible to spamming.

Constant monitoring is in part automated and in part administrator managed. System updates are performed daily. Monitoring includes: 1) Our servers are under constant packet-based and root-kit monitoring by the Duke University Health System security team, using a variety of enterprise tools. We also do similar monitoring using an independent set of tools. 2) We regularly monitor system memory, disk, runtime execution, and internet access profiles on the server through hand-written tools. This allows us to spot unusual activity as it occurs, and also to kill hung jobs, freeing capacity for other users. 3) The data directories are hand-scanned for Trojans on a regular basis, and are wiped several time a day. 4) We monitor the httpd and MolProbity logs for unusual activity and use firewall banning against suspicious URLs or those submitting batch runs too large for the server to handle. This too is accomplished with a set of hand-written scripts which are frequently modified as new threats emerge. We are committed to open, worldwide access for all types of legitimate users, so if your address seems to be incorrectly blocked, please send us email.

Unfortunately, it seems likely that serious, targeted attacks will be a growing problem for any scientific web site that performs active, open services for a worldwide user community. This puts a very significant burden on system management, in order to avoid exploit, denial-of-service or other mischief that could force closure of the site.

### II.4 Restoration of online kinemage graphics with NGL Viewer

Our Java-based KiNG display and modeling program (Davis et al. 2004; Chen et al. 2009) has historically served two separate functions. First, it provided interactive viewing, as a Java applet, of the validation results in 3D on the query model, seamlessly online with no download or installation needed. Second, when installed on the user’s computer, KiNG offers a very wide set of user-friendly modeling and analysis capabilities. The deprecation of the Java applet functionality still allows all the locally-installed usage of KiNG but has destroyed its online use as part of a MolProbity run.

In response to this problem, we have replaced the online viewing functionality of KiNG with a modified version of the Javascript NGL Viewer developed for the RCSB PDB (Rose et al. 2015). NGL Viewer gives excellent user perception of the 3D relationships and utilizes WebGL to provide blazingly fast interaction. In order to display kinemages in NGL Viewer, with the assistance of Alex Rose, we created a parser to translate the various kinemage objects (vectors, ribbons, balls, dots, etc.) into JavaScript objects that could be used in NGL Viewer. For ease of creating the initial interface, we used a demo GUI provided in the NGL Viewer code to display and control these translated kinemage objects. This GUI is currently capable of displaying and controlling all types of MolProbity validation markup (see Figure 5) and is being tested on the MolProbity beta site. This restores a user’s ability, online within the web browser, during the run, to explore the multi-criterion kinemage that shows local clustering and severity of all validation outliers on the 3D model.

**Figure 5.**
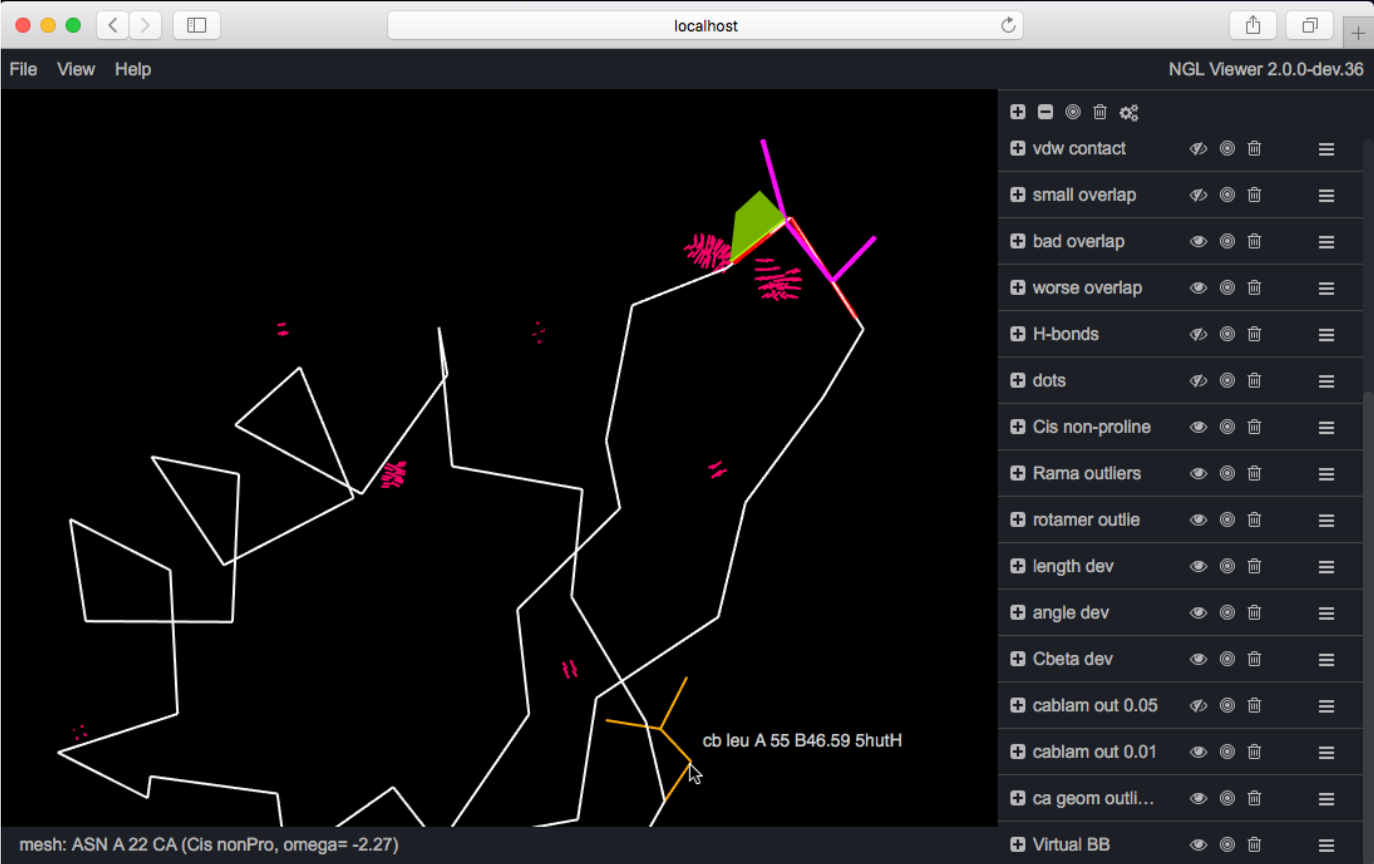
The demo GUI provided in the NGL Viewer code (Rose et al. 2015) has been modified to display and control kinemages of MolProbity validation markup. This screen capture shows a startup multi-criterion kinemage view with all validation flags on the Calpha trace.

The internal organization for the button-panel and mouse control for this GUI in NGL Viewer is quite different from the interface provided by KiNG and Mage (Richardson & Richardson 2001), so future plans for the software include a rewrite of the GUI. This rewrite will enable animation and better control over the kinemage groups and category “masters,” as well as a more streamlined interface.

### II.5 Expanded treatment of alternate conformation validation

Assessments that draw atoms from multiple residues present a particular challenge in comprehensive validation. For a given residue, Ramachandran analysis requires the C atom of the preceding residue in sequence for the calculation of phi, and the N atom of the succeeding residue for psi. Similarly, the calculations in CaBLAM require O and CA atoms across a span of 5 residues. Thus multiple alternate positions for an atom in an adjacent residue can result in multiple alternate validation results for a residue that contains no alternate positions itself.

Previous versions of MolProbity calculated and presented alternate validations only for the simple case of residues that themselves contained alternate positions. The new version calculates and presents all validations resulting from alternate atom positions. In the results table, some residues that do not contain alternate atom positions are now labeled as having alternates due to their proximity to residues that do. This more complete reporting of alternate conformations should guide users to a more complete understanding of alternates, especially at the sometimes problematic points at which alternate conformations rejoin non-alternate structure.

For clarity in defining and presenting percentages and residue counts, the overall statistics presented in MolProbity’s structure-level summary table are calculated from non-alternate residues plus alternate A only.

### II.6 Chirality checking

MolProbity has not had a check for chirality, because model-building libraries or fragments include only the correct forms, there are no chiral groups that span protein or nucleic acid monomer units, and standard refinement could not change chirality. However, we have recently encountered a few cases, including one where Amber refinement changed chirality at Cβ for a backward-fit Thr in order to pull all sidechain atoms into density peaks (Moriarty et al. 2019). We have now implemented a check of chiral volume outliers, which report only if there is a problem >4σ. It covers all the atoms defined as chiral centers in the Phenix Geo_std or monomer_library dictionaries. Three major cases are: 1) at Cα for D-amino-acids, either produced unintentionally or for genuine cases named as normal (e.g., Ala) rather than as D-amino-acids (e.g., Dal).; 2) at Cβ for Thr or Ile; 3) at substituents on RNA or DNA sugar rings. Some chiral-volume outliers are due to incorrect numbering in atom names, such as for OP1 vs OP2 on phosphates, or for numbering in Fe-S clusters, and we will either drop those or report them separately.

### II.7 New reference dataset in progress

High-resolution PDB deposits have more than doubled since our current Top8000 quality-filtered dataset of protein chains, and we are now set up to use map-density criteria for even better residue-level filtering (Hintze et al. 2016), so that we can obtain even cleaner data from about twice as many residues. Also, graphical databases are now available which are more suitable for our purposes than relational databases; we are in the process of switching to Neo4j (https://neo4j.com/docs/2.1.5/introduction.html).

At our present stage of database completion, we have already generated complete Cβ deviation plots as a function of amino-acid type and of χ1, which were not as exciting as we had hoped. We have also started to explore multi-dimensional ϕ,ψ,χ plots, which can differ even more dramatically by χ than by amino acid; Asn and Gln are shown in Richardson, Williams, and Richardson (2019). This new Neo4j database will soon be organized and populated enough to enable improved updates of the previous data distributions for all our conformational model validation criteria, especially valuable to improve the sensitivity level of CaBLAM outliers and occurrence statistics for UnDowser.

### II.8 Application of CaBLAM to cryoEM

An especially influential change in direction and emphasis since the previous Tools issue (Williams et al. 2018) has come mainly from our experience as assessors in two community exercises sponsored by the EMDB (Electron Microscopy Data Bank; https://www.ebi.ac.uk/pdbe/emdb/): the CryEM Model Challenge in 2016 (Richardson et al. 2018) and the CryoEM Model Metrics Challenge in 2019 (http://challenges.emdataresource.org/?q=model-metrics-challenge-2019). We also studied crystal structures at 3-4Å where a reliable answer was available at higher resolution.

That experience has demonstrated, in actual use, that CaBLAM supplies newly discriminating and robust overall and local validation information in the regime where traditional model validations fail: starting at 2.5Å to 3Å resolution and especially severe by 3.5 to 4Å, for either cryoEM or X-ray structures (Richardson et al. 2018b; Croll 2018). Significantly different models can fit equally well to those broad density maps. If traditional outliers do occur, they still indicate real problems, but at these resolutions they can be “gamed” simply by applying restraints that shove outliers across the nearest boundary, which seldom fixes and often worsens the underlying problems, so that scores make the models look much better than they actually are.

The recent cryoEM “resolution revolution” was enabled by direct electron detector hardware and by image collection as movies with software correction of specimen motion. It has resulted in dramatic numbers of new cryoEM structures at unprecedented resolutions better than 4Å, where *de novo* chain tracing is possible. Relative to x-ray crystal structures at these resolutions, refinement of models built into the cryoEM 3D reconstructions is done in real space rather than reciprocal space, phases are measured directly and start out quite good but are not improved by refinement, and backbone connectivity is typically somewhat better for cryoEM maps. The differences across a structure in effective resolution/disorder/uncertainty are even larger than for crystal structures. Since electrons are sensitive to local electrostatic potential, negatively charged carboxyls show weak density and Arg and Lys sidechains are strong. For nucleic acids, bases are strong and phosphates relatively weak although still fairly spherical and recognizable. Somewhere between 3.5 and 4Å resolution, nucleic acid double helices switch confusingly between connectivity across clear basepairs and connectivity along the direction of base stacking. For protein helices, in the range between 2.5 and 5Å resolution, the density gradually shifts confusingly between a spiral around a vacant axis and a tube with maximal density along the axis. In these transition regions some details in the density maps will give the wrong answer if followed too slavishly.

Relative to a density map at 2Å or better, 3 to 4Å density is broad, ambiguous, and sometimes even misleading. Therefore, model building and refinement must make use of more outside information to achieve physically reasonable models and well behaved refinement; even so, it is an extremely challenging task with extant tools. Covalent geometry (bond lengths and angles, planarity, etc.) needs to be quite tightly restrained. This causes no large problems because these are single-valued targets that cannot jump to some other allowed but incorrect local minimum. However, such tight restraints do destroy MolProbity’s Cβ deviation criterion Lovell et al., 2003), which flags incompatibility of sidechain and backbone conformation, such as a backward-fit branched sidechain. If geometry is perfect, then a Cβ cannot deviate from ideality even if perfect geometry puts it in the wrong position. In contrast, tight restraints on Ramachandran values or other multiple-minimum criteria make the scores better but the structure worse, since many of the changes pull the conformation into the wrong local minimum, as shown by examples below.

A major underlying problem is that at 2.5-3Å or worse, separate protrusions for the backbone carbonyl oxygens disappear into the tube of backbone density. Since those are the primary cue for fitting local backbone conformation, this makes mis-oriented peptides the commonest type of misfitting in 2.5 to 4Å cryoEM (or X-ray) structures. Whenever a peptide orientation is off significantly, the preceding ψ and following ϕ angles are grossly wrong. That almost always means that optimizing the Ramachandran values moves them into the wrong local minimum on the plot (Richardson et al. 2018b).

Fortunately, incompatibility of CO direction with the local Cα trace is the property most directly measured by CaBLAM. For each 5-residue stretch, it works from the relatively well-determined Cα virtual dihedrals to analysis of the much more dubious CO directions in the built model. (For more procedural detail, see Williams, Richardson, et al. in the 2018 Tools issue.) Very rare pairings of the Cα virtual dihedrals (plus the Cα virtual angle) are marked in red and provide an effective diagnosis of probable errors in Cα-only models, which are deposited fairly often near 4Å but have lacked suitable validation.

Since CaBLAM’s parameters span several residues and are not part of any current refinement program’s target set, its Cα-Cα-CO outliers can provide useful and independent validation at typical cryoEM resolutions. The 1% CaBLAM outliers are flagged graphically by magenta lines that follow the bad CO-CO dihedral, and on mouse-click or in numerical tables they provide conservative, quantitative diagnosis of which segments should be tried as regular α-helix or β-strand (Figure 6a, c). As true for other empirical-frequency validations, some outliers are genuine, and those are often of functional importance such as the ion-channel peptides in Figure 6b, in 6cju at 3.35Å (Rheinberger et al. 2018).

**Figure 6.**
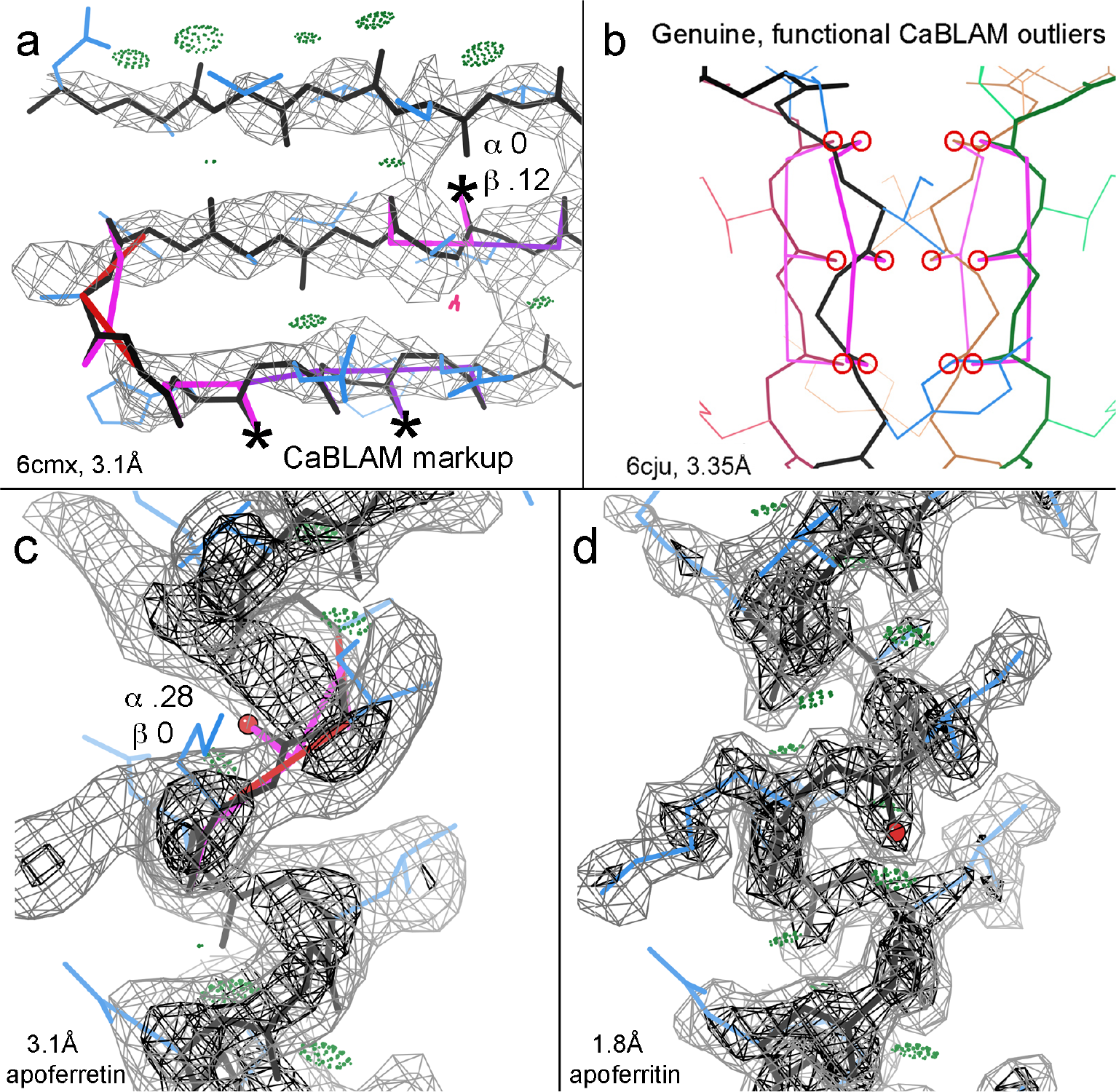
CaBLAM markup and secondary-structure diagnosis. a) CaBLAM suggests regular β-strands in 6cmx at 3.1Å (Li J et al. 2018), in spite of many outliers and few H-bonds. Each * CO just needs a near-180° peptide rotation. b) Genuine CaBLAM outliers form the ion-selectivity pore in 3.35Å 6cju (Rheinberger et al. 2018). c) CaBLAM suggests regular α-helix across a backward-pointing CO outlier in 3.1Å apoferritin density, and d) shows the CO correctly placed in its clear density at 1.8 Å.

CaBLAM markup for a β-sheet region is shown in Figure 6a, from the recent 6cmx cryoEM structure at 3.1Å resolution (Li J et al. 2018). The model is correctly trying to fit antiparallel β strands, but not successfully, because they make almost no β-type backbone H-bonds, and ϕ,ψ for 7/10 residues in the lower 2 strands are favorable but in the α or Lα Ramachandran regions, not in β. Here, CaBLAM 1% outliers flag a common lower-resolution pathology where 3 or more carbonyl O atoms point in the same direction rather than alternating, while the probabilities given for β conformation are in a highly suggestive range of 0.1 to 0.25. The central CO of such a triplet is almost always misoriented. For each of the COs marked with an asterisk, a near-opposite peptide rotation allows the CO to H-bond with an NH on a neighboring strand and removes the CaBLAM outliers. Once the two strands are corrected (not shown), the problematic turn can be refit as classic.

A very simple case of bad peptide orientation in α-helix is shown in Figure 6c. It comes from a model in the 2019 CryoEM Challenge, fit into a 3.1Å apoferritin map. The CO of Ile 145 is flanked by two CaBLAM outliers, it points nearly opposite the other carbonyls, the α probablity is 0.28, and preceding sidechains are pushed up from their density. The 1.8Å apoferritin map in Figure 6d shows the O density side peak clearly, with a well-fit model. If initial model-building starts with ideal secondary structures (which we would recommend at resolutions poorer than 2.5Å), then such outliers will not occur inside a helix or strand. However, outliers are still likely at the ends, and extending the regular α or β conformation should be tried as a likely correction. For instance, such a correctable C-terminal case (not shown) occurs at Lys 242 of 4heL at 3.2Å (Meena & Saxena, unpublished), and an N-terminal x-ray case at 3.5Å is shown and discussed in Moriarty et al. 2019.

Backbone conformations, CaBLAM outliers, and their corrections are much more diverse in turns and loops. A few are genuine, some are minor, and some flag major errors that may or may not show any other validation outliers. The probabilities given by CaBLAM for α or β conformation are seldom as high as 0.01. Figure 7a shows a CaBLAM outlier in the 99-104 loop of the 1de9 DNA-complex crystal structure at 3.1Å (Mol et al. 2000). Inspection shows residues out of density, which can be rebuilt (Figure 7b) once the problem is noticed.

**Figure 7.**
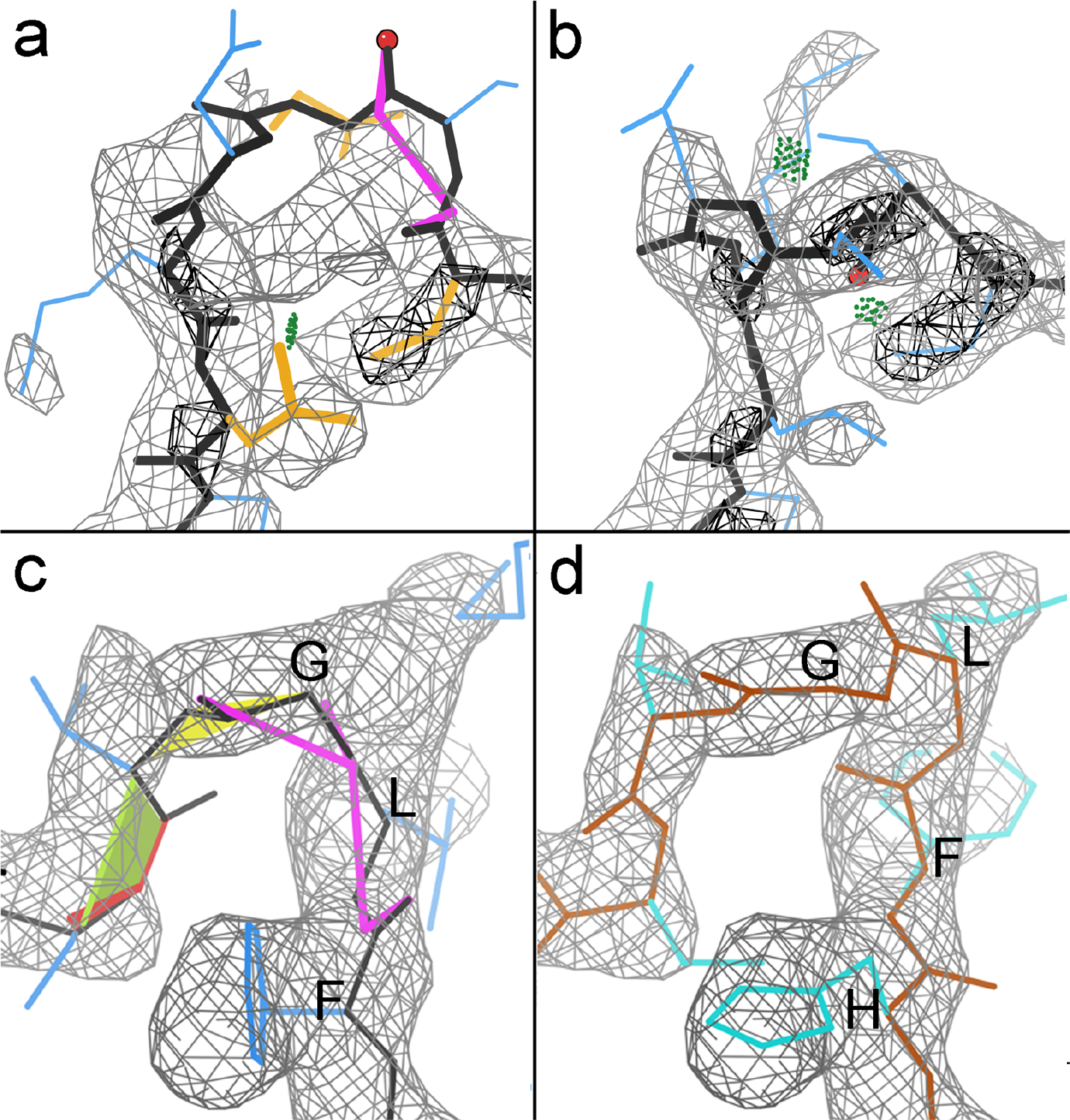
CABLAM markup as a guide to rebuilding. a) A CaBLAM outlier in the 99-104 loop of the 1de9 DNA-complex crystal structure at 3.1Å (Mol et al. 2000). b) The rebuilt region fitting into density. c) Indicates a missing residue by misalignment and poor fit of a loop in the alcohol dehydrogenase target at 2.9Å in the 2019 CryoEM Challenge, and d) shows a different, correct Challenge model with the sequence aligned correctly and no outliers.

If a model has a stretch of sequence misalignment, a disturbingly likely and serious problem at 3Å or worse, CaBLAM outliers very often flag one or both ends of the misaligned region. Figure 7c shows an example from the alcohol dehydrogenase target at 2.9Å in the 2019 CryoEM Challenge. A human alerted by multiple outliers can see that this model curves too tightly around the loop to fill its density, suggesting a missing residue. Indeed, Figure 7d shows a different, correct Challenge model which has the sequence aligned correctly and no outliers.

We and others are working toward automated correction for certain classes of CaBLAM outliers, or perhaps even initial avoidance of them. With guidance from the clear examples above at about 3Å, manual correction should be feasible in Coot (Emsley 2010) or Isolde (Croll 2018) for similar cases across the 2.5 to 4Å range. Start with what looks easiest: in or at the end of secondary structure, or two successive outliers in an otherwise good context.

## III. Discussion

We are very pleased to now have NGL Viewer as a replacement for online kinemage viewing, and UnDowser as a good initial utility for broad diagnosis of problematic peaks modeled as HOH. This new UnDowser tool, along with examples and descriptions for each scenario, should make it quick and easy for a structural biologist, or even an end-user, to come up with better reassignments for most clashing “waters” in a model. When implemented inside Phenix, it can become synergistic with existing tools for ion identification and other ligand identification, which have complementary strengths and shortcomings to those in UnDowser.

We have so far survived, at considerable cost, the highly targeted hacks on the MolProbity server. There is likely to become an increasingly serious need to develop and share protocols for providing open, worldwide access to important scientific web servers without a crippling level of vulnerability.

The recent cryoEM “revolution” is rapidly generating an unprecedented treasure of large, complex, dynamic, and biologically important structures. But, as explained here, there is a potentially dangerous disconnect in the process that could allow incorrect models and incorrect conclusions to go undetected until scientific contradictions pile up, as happened for similar reasons in crystallography around 1990 (see Introduction). The old measures developed since then (Rfree, Ramachandran, all-atom clashes, etc.) are still necessary but no longer sufficient, because the broad density at resolutions poorer than about 2.5Å is compatible with multiple quite distinct local models, both correct and incorrect. Therefore a new set of validation criteria are badly needed that are independent, not easy to refine directly, and diagnostic of local conformation but spanning more than a single residue to couple better with the larger-scale shapes in lower-resolution density. Our CaBLAM is almost the only current criterion with those properties, and it has indeed proven both sensitive and useful for making corrections, in practical use with cryoEM models. We and others will be working to develop further independent criteria that can collectively fill this validation gap.

## IV. Methods

The overall workflow, and the underlying methods, for the prior state of MolProbity were described in the previous Tools issue (Williams et al. 2018a) and have not changed.

UnDowser runs automatically along with the clashscore assessment, as it uses the same Reduce’d file and calculations but adds water-water interactions. Its input is the set of HOH entities in the coordinate file that have at least one all-atom clash (non H-bond overlap ≥ 0.4Å). Its output table is modeled on the MolProbity multi-criterion chart, with a row for each clash interaction, sorted first by total severity for each water (Sum(overlap −0.2Å), and within each water by individual clash severity. The row groups for each water are distinguished by alternate pale gray or white coloring. After complete identifiers and B-factors for each atom in the clash, succeeding columns classify each clash by the type of atom the HOH clashes with (polar, nonpolar, other water, or alternate-conformation atom), with clash severity color-coded progressively as pink, hotpink, or red by the same divisions of clash overlap (0.4, 0.5, 0.9) as used in the main MolProbity multi-criterion chart. The assignment of possible interpretation(s) for consideration and inspection are based on the characteristics of manually-evaluated examples such as those shown in the text. For instance, for an HOH that has ≥2 clashes with full or partial negative charges, no clashes with nonpolars, no clashes or H-bonds with positive polars, a B-factor close to or less than the average of surrounding atoms, and no alternate conformations involved, a positive ion will be the major suggestion, with a note to check for very low density of the HOH. For an HOH that clashes with the first or last modeled backbone atom at a non-terminal chain end, the major suggestion is to try extending the chain further, starting with that clashing HOH. The rules are preliminary and will be reformulated after large-scale runs with the new reference dataset and their analysis.

The chirality check is run automatically along with covalent geometry validation, since it works from the same file produced by mmtbx.mp_geo. Chiral volume outliers >10σ are noted as likely changes of handedness.

The modified Javascript code to show multi-criterion kinemages in NGL Viewer is available on GitHub in a fork from the main NGL Viewer repository, at https://github.com/vbchen/ngl.

The methods behind CaBLAM are described in the previous Tools issue (Williams et al. 2018b). Both the Python code and the underlying reference-data distributions are available from the rlabduke repository on GitHub, and are also distributed with Phenix (Liebschner et al 2019). Most CaBLAM examples and conclusions here were drawn from our assessments at the 2016-17 EMDB CryoEM Model Challenge (Richardson et al. 2018b) and at the 2019 EMDR Model Metrics Challenge (not yet published). In 2019 we used both our own MolProbity runs, including CaBLAM, and the scores, superpositions, and comparisons made available by Andriy Kryshtafovych on the Challenge web site (http://challenges.emdataresource.org/?q=model-metrics-challenge-2019), on all submitted models for all 4 targets. Reference models (assumed as essentially correct) were 3ajo (Masuda et al. 2010) for 1.8Å, 2.3Å, and 3.1Å effective resolutions of apoferritin, and 6nbb (Herzik et al. 2019) for the alcohol dehydrogenase dimer target at 2.9Å. We visualized CaBLAM outliers and density maps on multi-criterion kinemages in KiNG (Chen et al. 2009) and analyzed their deleterious effect on ϕ,ψ values in kinemage Ramachandran plots with clickable and searchable residue datapoints. Outlier correction was defined on the Challenge targets by the reference-structure conformation, and for other structures the outliers were manually corrected in KiNG or Coot, with success declared if the result had no outliers and same or better map fit.

All figures except 1 and 5 were made in KiNG. PDB codes are given as lowercase except for L (4heL,1qLw), a convention that gives no ambiguities in any font, such as 1/I/l or O/0.

## References

Arendall BW III, Tempel W, Richardson JS, Zhou W, Wang S, et al. (2005) A test of enhancing model accuracy in high-throughput crystallography, J. Struc. Func. Genomics 6: 1011

Banuelos S, Saraste M, Carugo KD (1998) Structural comparisons of calponin homology domains: implications for actin binding, Structure 6: 1419–1431 [1bkr]

Barad BA, Echols N, Wang RY-R, Cheng YC, DiMaio F, Adams PD, Fraser JS (2015) EMRinger: side-chain-directed model and map validation for 3D electron microscopy, Nat. Meth. 12: 943–946

Bartesaghi A, Merk A, Banerjee S, Matthies D, Wu X, Milne J, Subramaniam S (2015) 2.2Å resolution cryo-EM structure of beta-galactosidase in complex with a cell-permeant inhibitor, Science 348: 1147–1151 [5a1a]

Berman H, Henrick K, Nakamura H (2003) Announcing the worldwide Protein Data Bank, Nat. Struct. Mol. Biol. 10: 980

Borell B (2009) Fraud rocks protein community, Nature 462: 970

Bourne PC, Isupov MN, Littlechild JA (2000) The atomic resolution structure of a novel bacterial esterase, Structure 8: 143–151 [1qLw]

Brunger AT (1988) Crystallographic refinement by simulated annealing: Application to a 2.8Å resolution structure of aspartate aminotransferase, J. Mol. Biol. 203: 803–816

Brunger AT (1992) The free R value: A novel statistical quantity for assessing the accuracy of crystal structures, Nature 355: 472–474

Chen VB, Davis IW, Richardson DC (2009). “KiNG (Kinemage, Next Generation): A versatile interactive molecular and scientific visualization program”, Protein Sci 18: 2403–2409

Chen VB, Arendall WB III, Headd JJ, Keedy DA, Immormino RM, et al. (2010) MolProbity: all-atom structure validation for macromolecular crystallography, Acta Cryst. D66:12–21

Croll TI (2015) The rate of *cis-trans* conformational errors is increasing in low-resolution crystal structures, Acta Cryst. D71: 706–709

Croll TI (2018) ISOLDE: a physically realistic environment for model building into low-resolution electron-density maps, Acta Cryst. D74: 519–530 [6eyc]

Davis IW, Murray LW, Richardson JS, Richardson DC.(2004) MolProbity: structure validation and all-atom contact analysis for nucleic acids and their complexes, Nucleic Acids Res. 32: W615–W619

Davis IW, Arendall WB III, Richardson JS, Richardson DC (2006) The backrub motion: How protein backbone shrugs when a sidechain dances, Structure 14: 265–274

Davis IW, Leaver-Fay A, Chen VB, Block JN, Kapral GJ, et al. (2007) MolProbity: All-atom contacts and structure validation for proteins and nucleic acids, Nucleic Acids Res. 35: W375–W383

Echols N, Morshed N, Afonine PV, McCoy AJ, Miller MD, et al. (2014) Automated identification of elemental ions in macromolecular crystal structures, Acta Cryst. D70: 1104–1114

Emsley P, Lohkamp B, Scott WG, Cowtan K (2010) Features and development of Coot, Acta Cryst. D66: 486–501

Gore S, Velankar S, Kleywegt GJ (2012) Implementing an X-ray validation pipeline for the Protein Data Bank, Acta Cryst. D68: 478–483

Hayashi I, Oda T, Sato M, Fuchigami S (2018) Cooperative DNA binding of the plasmid partitioning protein TubR from the *Bacillus cereus* pXO1 plasmid, J. Mol. Biol. 430: 5015–5028 [6aht]

Headd J, Richardson J (2013) Fitting Tips #5: What’s with water?, Comput. Cryst. Newsletter 4: 2–5

Herzik MA Jr, Wu M, Lander GC (2019) High-resolution structure determination of sub-100 kDa complexes using conventional cryoEM, Nat. Commun. 10: 1032–1032 [6nbb]

Hintze BJ, Lewis SM, Richardson JS, Richardson DC (2016) MolProbity’s ultimate rotamer-library distributions for model validation, Proteins: Struc Func Bioinf 84: 1177–1189

Hooft RWW, Vriend G, Sander C, Abola EE (1996) Errors in protein structures. Nature 381: 272

Hwang KY, Song MJ, Kim JS, Cheong WC (2019) unpublished [6a4v]

Janssen BJC, Read RJ, Brunger AT, Gros P (2007) Crystallography: crystallographic evidence for deviating C3b structure, Nature 448: E1–E3

Jones TA, Zou JY, Cwan SW, Kjeldgaard M (1991) Improved methods for building protein models in electron density maps and the location of errors in these models, Acta Cryst. A47: 110–119

Knight S, Andersson I, Branden C-I (1989) Reexamination of the three-dimensional structure of the small subunit of RuBisCo from higher plants, Science 244: 702–705

Kuratani M, Hirano M, Goto-Ito S, Itoh Y, Hikida Y, Nishimoto M et al. (2010 Crystal structure of Methanocaldococcus jannaschii Trm4 complexed with sinefungin, J. Mol. Biol. 401: 323–333 [3ajd]

Laskowski RA, MacArthur MW, Moss DS, Thornton JM (1993) PROCHECK: a program to check the stereochemical quality of protein structures. J. Appl. Crystallogr. 26: 283–291

Li S, Finley J, Liu Z-J, Qiu SH, Luan CH, et al. (2002) Crystal structure of the cytoskeleton-associated protein glycine-rich (CAP-Gly) domain, J. Biol. Chem. 277: 48596–48601 [1LpL]

Li H, Zhang W, Dong C (2018) Crystal structure of the outer membrane protein OmpU from *Vibrio cholerae* at 2.2Å resolution, Acta Cryst. D74: 21–29 [5onu]

Li J, Shalev-Benami M, Sando R, Jiang X, Kibrom A, et al. (2018) tructural basi for teneurin function in circuit-wiring: A toxin motif at the synapse, Cell 173: 735–748.e15 [6cmx]

Liebschner D. Afonine PV, Baker ML, Bunkoczi G, Chen VB, et al. (2019) Macromolecular structure determination using X-rays, neutrons, and electrons: Recent developments in Phenix, Acta Cryst D in press

Lovell SC, Davis IW, Arendall WB III, de Bakker PIW, Word JM, Prisant MG, Richardson JS, Richardson DC (2003) Structure Validation by Cα Geometry: ϕ,ψ and Cβ Deviation, Proteins: Struct Funct Genet 50: 437–450

Macauley KE, Jia-Xing Y, Dodson EJ, Lehmbeck J, Ostegaard PR, Wilson KS (2001) A quick solution: *Ab initio* structure determination of a 19 kDa metalloproteinase using Acorn, Acta Cryst. D57: 1571–1578 [1eb6]

Masuda T, Goto F, Yoshihara T, Mikami B (2010) The universal mechanism for iron translocation to the ferroxidase site in ferritin, which is mediated by the well conserved transit site, Biochem. Biophys. Res. Commun. 400: 94–99 [3ajo]

Meshcheryakov VA, Krieger I, Kostyukova AS, Samatey FA (2011) Structure of a tropomyosin N-terminal fragment at 0.98Å resolution, Acta Cryst. D67: 822–825 [3adv]

Montelione GT, Nilges M, Bax A, Guntert P, Hermann T, et al. (2013) Recommendations of the NMR structure validation task force, Structure 21: 1563–1570

Mol CD, Izumi T, Mitra S, Tainer JA (2000) DNA-bound structures and mutants reveal abasic DNA binding by APE1 and DNA repair coordination, Nature 403: 451–456

Moriarty NW, Janowski PA, Swails JM, Nguyen H, Richardson JS, Case DA, Adams PD (2019) Improved chemistry restraints for crystallographic refinement by integrating Amber molecular mechanics into Phenix, Acta Cryst. D (submitted) and bioRxiv, 724567

Murshudov GN, Grebenko AI, Brannigan JA, Antson AA, Barynin VV, Dodson GG, Dauter Z, Wilson KS, Melik-Adamyan WR (2002) The structure of *Micrococcus lysodeikticus* catalase, its ferryl intermediate (compound II) and Nadph complex, Acta Cryst. D58: 1972–1982 [1gwe]

Pai EF, Krengel U, Petsko GA, Goody RS, Kabsch W, Wittinghofer A (1990) Refined crystal structure of the triphosphate conformation of H-ras p21 at 1.35Å resolution: implications for the mechanism of GTP hydrolysis, EMBO J. 9: 2351–2359

Pintile G, Chiu W (2018) Asessment of structural features in cryo-EM density maps using SSE and sidechain Z-scores, J. Struct. Biol. 204: 564–571

Read RJ, Adams PD, Arendall WB III, Brunger AT, Emsley P, et al. (2011) “A New Generation of Crystallographic Validation Tools for the Protein Data Bank”, Structure 19:1395–1412

Reisky L, Prechoux A, Zuhlke MK, Baumgen M, Robb CS, et al. (2019) A marine bacterial enzymatic cascade degrades the algal polysaccharide ulvan, Nat. Chem. Biol. 15: 803–812 [6hhm]

Rheinberger J, Gao X, Schmidpeter PA, Nimigean CM (2018) Ligand discrimination and gating in cyclic nucleotide-gated ion channels from apo and partial agonist-bound cryo-EM structures, Elife 7[6cju]

Richardson DC, Richardson JS (2001) Mage, Probe, and Kinemages, chapter 25.2.8 in IUCr’s *International Tables of Crystallography*, Volume F: Crystallography of Biological Macromolecules (Eds: Michael G. Rossmann and Eddy Arnold). Kluwer Academic Press: Dortrecht

Richardson JS, Richardson DC, Tweedy NB, Gernert KM, Quinn TP, et al. (1992) Looking at proteins: representations, folding, packing, and design, Biophys. J. 63: 1186–1209

Richardson JS, Richardson DC (2013) Doing molecular biophysics: Finding, naming, and picturing signal within complexity, Ann. Rev. Biophysics 42:1–28

Richardson JS, Williams CJ, Hintze BJ, Chen VB, Prisant MG, Videau LL, Richardson DC (2018a) “Model validation -- local diagnosis, correction, and when to quit”, Acta Cryst D74: 132–142

Richardson JS, Williams CJ, Videau LL, Chen VB, Richardson DC (2018b) Assessment of detailed conformations suggests strategies for improving cryoEM models: helix at lower resolution, ensembles, pre-refinement fixups, and validation at a multi-residue length scale”, J Struct Biol 204: 301–312

Richardson J, Richardson D, Williams C (2019) Fitting Tip #17 - Asn and Gln are remarkably different, Comp. Cryst. Newsletter 10: 1–6

Rose AS, Hildebrand PW (2015) NGL viewer: a web application for molecular visualization, Nucleic Acids Research 43: W576–W579

Williams CJ, Richardson JS (2015) Fitting Tips #9: Avoid excess *cis* peptides at low resolution or high B, Comp Cryst Newsletter 6: 2–6

Williams CJ, Hintze BJ, Headd JJ, Moriarty NW, Chen VB, et al. (2018a) MolProbity: More and better reference data for improved all-atom structure validation, Protein Science 27: 293–315

Williams CJ, Videau LL, Hintze BJ, Richardson JS, Richardson DC (2018b) “*Cis*-nonPro peptides: Genuine occurrences and their functional roles”, bioRxiv, 324517

Winn MD, Ballard CC, Cowtan KD, Dodson EJ, Emsley P, et al. (2011) Overview of the CCP4 suite and current developments, Acta Cryst. D67: 235–242

Wlodawer A, Miller M, Jaskolski M, Sathyanarayana BK, Baldwin E, et al. (1989) Conserved folding in retroviral proteases: crystal structure of a synthetic HIV-1 protease, Science 245: 616–621

Word JM, Lovell SC, LaBean TH, Zalis ME, Presley BK, Richardson JS, Richardson DC (1999a) Visualizing and quantitating molecular goodness-of-fit: Small-probe contact dots with explicit hydrogen atoms, J Mol Biol 285: 1711–1733

Word JM, Lovell SC, Richardson JS, Richardson DC (1999b) Asparagine and glutamine: Using hydrogen atom contacts in the choice of side-chain amide orientation, J Mol Biol, 285: 1735–1747

Zheng H, Cooper DR, Porebski PJ, Habalin IG, Handing KB, Minor W (2017) CheckMyMetal: a macromolecular metal-binding validation tool, Acta Cryst. D73: 223–233

